# Designer DNA Strand Displacement Reaction toward Controlled Release of Cargos

**DOI:** 10.1101/2024.07.23.604118

**Authors:** Chih-Hsiang Hu, Remi Veneziano

## Abstract

Dynamic DNA nanotechnology systems are used to design DNA logic circuits, signal amplification mechanisms for biosensing, and smart release system that could potentially be used in several biomedical applications. The toehold-mediated strand displacement reaction (**TMSDR**) is one of the main methods for designing DNA-based biomolecular logic circuits. However, the reaction behaviour such as the displacement rate and the quantity of strand released are difficult to control and often requires chemically modified strands or addition of enzymes. This makes the TMSDR versatility and specificity limited, and not always adapted for biomedical applications. Therefore, further understanding the sequence design parameters enabling fine tuning of the TMSDR behaviour without the need for complex modification, would enable its broader application. In this study, using a DNA motif developed for multiplexed release, we examine how mismatched base(s) in the trigger strand is affecting the release rate and quantity released and found that both location and type of mismatched base(s) significantly impact the displacement parameters of the TMSDR. This allows for a finer control of the cargo release for the multiplexed release system that could be used for varying biomedical applications and help developing release system mimicking the natural distribution of biomolecules.

## INTRODUCTION

Dynamic DNA nanotechnology systems are powerful tools for designing a wide array of stimuli-responsive circuits for many biomedical and non-biomedical applications such as drug delivery system, biosensor, and computational system (1–6). These dynamic systems are of particular interest for applications that require precise control over both temporal release and distributions of biomolecules as they could mimic the biological signal found in tissues and enables the development of smart biomaterials for regenerative medicine. Thus, thanks to the nature of DNA and the variety of DNA motifs available, there are many stimuli that have already been used to generate a dynamic DNA nanotechnology system, which include pH, light, temperature, enzymatic reactions, strand displacement, and aptamer (7–12). Amongst these stimuli for designing dynamic DNA nanotechnology system, the toehold-mediated strand displacement reaction (**TMSDR**), which has been used for DNA circuits (13–17), biosensors (4, 5, 18), release system (19, 20), reconfigurable nanodevices (21–23) is particularly attractive. Indeed, the TMSDR operates by using a single strand DNA (trigger strand) with higher affinity to displace another strand in a DNA duplex (incumbent) and does not require any other external stimuli (**Figure S1**). This reaction is achieved by having a toehold region, a single strand DNA overhang, on the binding strand that leads to the higher affinity to the trigger strand compared to the incumbent strand. There have been various studies on understanding how TMSDR works and how to control the behaviour of the TMSDR system such as displacement rate to design specific dynamic system. One of the main methods to modulate the behaviour of the TMSDR system is to incorporate chemically modified DNA sequence or additional components, such as enzymes or modulating the DNA concentration (24–27). However, those techniques are not always suitable for biomedical applications due to potential biocompatibility issues caused by these modifications or by the absence of such enzymes in the targeted tissues. Additionally, instead of chemically modifying DNA or adding components, the ability to modulate the TMSDR system by simply altar the DNA sequences used would offer more versatility and reduce the design time by preventing a complete redesign of the system. On the other hand, researchers also have been exploring other design parameters to control the behaviour of TMSDR system without the addition or modification of components. One of the main design parameters is the length of toehold region and it has been found to greatly affect the displacement rate of the system by order of magnitudes (28, 29). In additional to the toehold length, the presence of mismatched base(s) has also been studied as a potential design parameter to control the TMSDR behaviour (30–33). A notable design parameter with mismatches in TMSDR is the location of mismatch in the system and it has been shown that it will affect the displacement rate by orders of magnitudes depending on the overall TMSDR system design (32). On top of controlling the rate of TMSDR, researchers also have incorporated mismatches into DNA circuits to improve its performance and reduce leakage (30, 34, 35). However, none of the studies listed above have studied in detail the effect of these design parameters to define a set of rules that can be used to control the reactions rate. In this study we used a previously designed DNA nanostructure in a form of a 4-way junction (**4WJ**) presenting up to three strand displacement systems was utilized instead of the common DNA duplex toward designing of a sequential release system that would be able to deliver multiple biological molecules on demand. Specifically, we aim to further understand how the displacement rate and quantity of a TMSDR system is affected by sequence design, specifically the incorporation of mismatched bases in the trigger strand. To achieve that goal, we designed various trigger sequences with different mismatch conditions (i.e., location, quantity, and types of mismatches) and examine how those conditions affect the displacement of our system. Furthermore, based on some of our experimental results and previous studies, we developed a model that could potentially predict the displacement properties based on the mismatch condition presented.

## MATERIAL AND METHODS

### Material

All oligonucleotides, modified and unmodified, (Table S1, S2, S3) were purchased from Integrated DNA Technologies (Coralville, IA, USA) and used without further purification. Water, molecular biology grade (351-029-101) was purchased from Quality Biological (Gaithersburg, MD). 10X PBS (J75889) was purchased from Thermo Scientific (Waltham, MA).

### Trigger sequence Design

#### Trigger with Varying Toehold Length

Triggers were designed with varying toehold length (3 to 7 bases) to establish baseline for our system and compared with the well-established literature to ensure our system is behaving similarly to the established systems (21, 23, 28, 29).

#### Trigger with Mismatched Bases

The triggers with mismatched base(s) were designed based on three parameters: the mismatch location, mismatch quantity, and mismatch type. The mismatch location describes the location (from 5’ to 3’) where the mismatch base(s) occurs. The mismatch quantity describes the total number of mismatch bases present in the trigger strand. The mismatch type describes the mismatch base present on the trigger strand and compared to corresponding hybridizing strand (target strand); therefore, with the 4 bases, there are a total of 12 mismatch types possible that we identify using a numerical representation for simplicity (Table 1). Trigger design schematic can be found in SI (**Figure S2**).

**Table 1:** All possible mismatch types and its numerical representation for this study. Mismatch type is shown as the ratio of the mismatch base on the trigger strand:corresponding base on the target strand. The group number is based on which original base the mismatch replaced.

| Group Number | Numerical Representation | Mismatch Type |
| --- | --- | --- |
| 1 | 1 | T:T |
|  | 2 | G:T |
|  | 3 | C:T |
| 2 | 4 | A:A |
|  | 5 | C:A |
|  | 6 | G:A |
| 3 | 7 | A:G |
|  | 8 | T:G |
|  | 9 | G:G |
| 4 | 10 | A:C |
|  | 11 | T:C |
|  | 12 | C:C |

### Experimental methods

#### DNA Motif Folding

The 4-way junction (**4WJ**) DNA motif is folded as described in our previous work (19). Briefly, the DNA strands were mixed equimolarly at 5 µM in 1X folding buffer (40 mM TRIS-base, 20 mM acetic acid, 2 mM EDTA, 12 mM MgCl_2_, at pH 8.0) and annealed in a Bio-Rad T100^TM^ Thermal Cycler using a temperature ramp from 95 to 4℃ over the course of 2h. DNA strands modified with fluorophore (HEX or Texas Red [**TR**]) or quencher (Iowa Black FQ or Iowa Black RQ) were incorporated into the overhang of the 4WJ. The DNA sequences used for the 4WJ folding can be found in SI (**Table S1**).

#### DNA Motif Characterization

Agarose gel electrophoresis was used to characterize the 4WJ DNA motif to ensure the structure was folded properly using the established method in our previous paper (19). Briefly, 1.5 wt% high melt agarose gel was made in Tris Acetate EDTA (TAE) and 30 µL of DNA motifs at 250 nM in 1X folding buffer were loaded and ran at 110 V for 30 minutes. The QuickLoad 100 bp DNA ladder from New England BioLabs® (Ipswich, MA, USA) was used along with the samples.

The hydrodynamic diameter of the 4WJ motif was measured using dynamic light scattering (DLS) similarly with our previous study (19). The 60 µL of 750 nM 4WJ was prepared with 1X PBS and used to measure the hydrodynamic diameter of the 4WJ.

#### Spectroscopy Study for the Strand Displacement Reaction

Fluorescence of the DNA motif was measured for fluorophore (HEX or TR) samples, quenched (HEX/Iowa Black FQ or TR/Iowa Black RQ) samples, and the unmodified samples using the established method in our previous paper (19). Briefly, the DNA motifs were diluted to 500 nM for all conditions. For the HEX fluorophore (Ex: 540 nm, Em: 570 nm) pair, it was excited at 500 nm and recorded from 535 to 800 nm. For the TR fluorophore (Ex: 585 nm, Em: 618 nm) pair, it was excited at 565 nm and the emission spectrum was recorded from 590 to 800 nm. The quenching efficiency of the quenchers was calculated based on Equation (1).

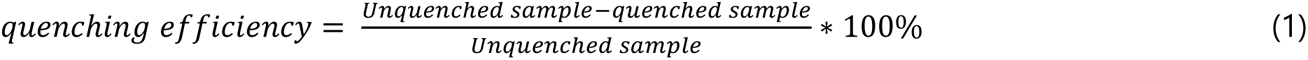

The spectroscopy study in qPCR was conducted as descripted in our previous work (19). Briefly, the fluorescence was measured over the course of ∼3 hours at 40℃ using the “Q” qPCR machine from Quantabio. All studies were carried out in 1X PBS and 0.1X folding buffer condition with 500 nM of 4WJ motif. HEX fluorophore pair were used as strand displacement probe system and the fluorescence results was measured in yellow channel (λ = 570 nm). The control samples without trigger and without quencher were used for negative and positive control, respectively. There are three main types of trigger experimental condition tested in the spectroscopy study: concentration of the trigger, trigger design (mismatch bases), and ratio of trigger:4WJ. For all three study types, displacement behaviour (*i.e.,* rate and binding quantity) were examined. For trigger concentration, the concentration of the trigger strands was tested between 0 to 2500 nM (0-5X molar ratio trigger: 4WJ) at different increments. For trigger design, the concentration of the trigger strands is fixed at 500 nM, which corresponded to 1:1 ratio between the 4WJ motif and the trigger. For the trigger ratio, the trigger mixture is composed with two different triggers and the total trigger concentration of the trigger mixture is fixed at 500 nM, which corresponded to 1:1 ratio between the 4WJ motif and trigger; however, the ratio of the trigger strands was varied within the mixture.

In addition to the HEX fluorophore, TR fluorophore and Iowa Black quencher were incorporated at a different overhang on the 4WJ. TR/Iowa Black pair was also able to undergo strand displacement reaction using similarly designed trigger as the HEX/Iowa Black pair. The result was collected from the red channels (λ = 618 nm) for TR instead of the yellow channel. Similar experimental procedure described above was carried out for the Texas Red/Iowa Black pair and the results were compared to the predicted result based on the result of the HEX/Iowa Black pair trigger screening. Additionally, both HEX/Iowa Black and TR/Iowa Black pairs were incorporated into the 4WJ and triggered with two different triggers to demonstrate the feasibility of our system for multiplex controlled release application.

Furthermore, a mixture of triggers was used to study the multiplex release system. The mixture of triggers composed of one trigger for HEX fluorophore and one trigger for TR fluorophore. The individual trigger:4WJ ratio was 1:1 and the fluorescence signals were recorded in two separate channels on the ‘Q’ qPCR machine simultaneously. The final trigger:4WJ ratio is 2:1 since there are two different triggers presented in the system.

#### Stability assessment

The stability of the DNA motif at testing conditions was tracked by incorporating the control samples (4WJ without trigger and 4WJ without quencher) in our study at identical buffer and temperature conditions. The fluorescence of both 4WJ without trigger and 4WJ without quencher was recorded over the total experimental time (∼2 hours).

### Data Analysis

#### Raw Data Processing

The data files (.csv) from the fluorescence displacement studies were processed using the Python3.xx with *Panda* package. The general data processing involves the subtraction of average negative control, normalization against the full complementary trigger result. This process is carried out for each independent data file. After the normalization step, the results were combined based on the trigger strand used in the assay into one dataframe and summarized with average and standard deviation at each time point.

#### Model Fitting

The processed data underwent curve fitting using the *lmfit* package in Python with one-phase association with lag and second order kinetic models. Curve fitting for the one-phase association with lag was carried out using equation (2)

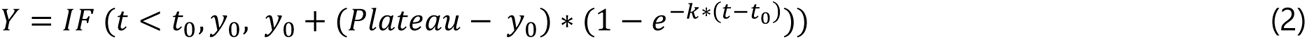

Where ’t_0_’ is when the reaction starts, ‘y_0_’ is the initial value prior ‘t_0_’, ‘Y’ is the release at a specific time ‘t’, ‘k’ is the rate constant with unit of time^-1^. The curve fitting program will fit ‘y_0_’, Plateau, and ‘k’ for all cases and if the reaction was not started at the start of the data recording, the ‘t_0_’ term will also be fitted to generate a more representative fit. However, if there was no initial lag observed, equation **Error! Reference source not found.**(2) was simplified into a one-phase association equation (3). The terms ‘k’, ‘y_0_’, and ‘t_0_’ were determined analytically using non-linear regression using least squared method.

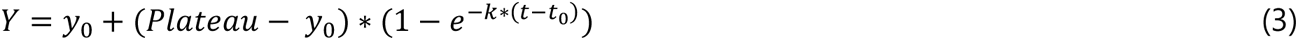

For second order kinetic model [equation (4)], simple ODEs were solved using *SciPy* and fitted to the experimental data using the *lmfit* package. For the second order kinetic model, the system was simplified without the reversible intermediate step and the reaction was assumed to be irreversible (30, 32).

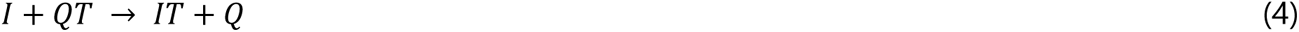

Where I, is the invader strand, QT is the initial complex, IT is the end complex, and Q is the quencher strand. The fitting was carried out to both overall and individual release profile. For the overall release profile fitting, a weighted curve fitting procedure was carried out using the average value and weighted using the inverse of calculated variance to generate the 3σ confidence band. For the individual fits, the standard curve fitting procedure was carried out using the release value from each individual trials. The r-squared value is recorded for all fitting procedures.

#### Displacement Behaviour Prediction via Linear Regression

The mismatch type and location from the HEX/Iowa Black pair was parameterized via dummy encoding to represent the presence of specific type or the location of the mismatch. The dummy variables are used to carry out regression against the plateau and rate value obtained from the one-phase association model. The perfectly matched trigger was omitted in the parameterisation process to avoid perfect collinearity of the dummy variable matrix and is represented as the intercept of the regression model. The regression model was then tested against the results obtained from TR/Iowa Black pairs.

#### Nupack Simulation

*Nupack v4.0.1.8* (36) was used as a package with Python3 to simulate the theoretical concentration for each component in the system at specific ion concentrations. The monovalent ion (sodium “Na^+^” and potassium “K^+^” ions) concentration is set to 161.5 mM and divalent ion (magnesium ions “Mg^2+^”) concentration is set to 1.2 mM. The different ion concentrations were calculated based on the buffer condition used in our experiments (1X PBS + 0.1X folding buffer). The temperature was set to be 40℃ for the Nupack model to be consistent with the experimental condition. The stacking option of the ensemble was selected for the Nupack model. The tube class was defined with three DNA strands (base, incumbent, and trigger) at the starting concentration of 500 nM. The maximum complexes size was set to be 3 for the tube class. Both complex analysis and complex concentration function from *Nupack* with the tube class and the model specified were used to calculate the structure free energy and simulate the theoretical concentration of each complex.

### Statistical Analysis

The statistical analyses were carried out in Python using the *SciPy*, *statsmodels*, and *pingouin* packages. Welch one-way analysis of variance (ANOVA) with Games-Howell post-hoc analysis was used to circumvent the potential issue arise from the uneven sample size and variance. Additionally, the effect size is calculated using eta-squared (η^2^) for ANOVA analysis to estimate how much the variant of the dependent variable is accounted by the independent variable (37, 38). For two-sample T-test, the unequal variance assumption was used and the effect size is calculated using Hedge’s g (g). Both Welch ANOVA analysis with Games-Howell post-hoc analysis and two-sample T-test for rate are calculated based on the log-transformed data. The data for each condition is collected in triplicate with four replicates for each trial. The regression analysis was carried out for each individual trial (n≥12) and the statistical analysis was carried out for all regression result that have a r^2^ > 0.5.

For the box plot presented in this paper, Each dot in the boxplot represents individual measurement. The whisker was set to be the 25^th^ and 75^th^ percentile of the median. The cross represents the potential outlier presented in the dataset.

The source code for the data analysis described above is available at GitHub (https://github.com/seanhu0401/DNA-Strand-Displacement-System.git).

## RESULTS & DISCUSSION

### Design, assembly, and characterization of the strand displacement system

A 4WJ motifs (Figure 1A) that was developed in a previous study for sequential strand displacement reactions to release multiple cargos was used in this study(19). After annealing of the 4WJ, DLS was carried out and the measured hydrodynamic diameter measured was (13.08 ± 2.04 nm) which is slightly smaller to the estimated diameter of 16.32 nm and similar to the measured diameter of 4WJ (13.45 nm) in our previous paper (19). A representative volume curve for the DLS measurement is shown in Figure 1B. Agarose gel electrophoresis was carried out to ensure 4WJ was folded properly with minimal by-products present. As shown in Figure 1C, the folding of the 4WJ appear to be complete with minimal by-products observed in the agarose gel. Next, we assembled the 4WJ with both fluorophore and quenchers to validate the formation of the strand displacement circuit. We first tested the quenching of the fluorophore used in presence of the quencher to ensure that a sufficient quenching (>95%) occurred for both fluorophores. We found that both HEX and TR were ∼99% quenched at the concentration measured (Figure 1D, E). The large quenching efficiency is required to ensure that the signals observed in our experimentations are from the release of quencher and not from non-specific emission, and particularly for multiplexing. There is a small fluorescent signal recorded with TR samples when using HEX excitation parameters due to the small overlap in the excitation spectra of both TR and HEX. However, this should not affect the TR measurement since it only contributes ∼5% of the real signal. Furthermore, in the spectroscopy study for the strand displacement reaction, both negative (**NT**) and positive (**NQ**) controls were used and both NT and NQ signals remain constant throughout the experimentation duration (∼2 hours) (Figure 1F), which indicates the 4WJ was stable for the experimental time and condition. Therefore, the increase in the fluorescence that will be further observed in the study will not be due to the degradation of the 4WJ and there is no release without the addition of trigger strand.

**Figure 1:**
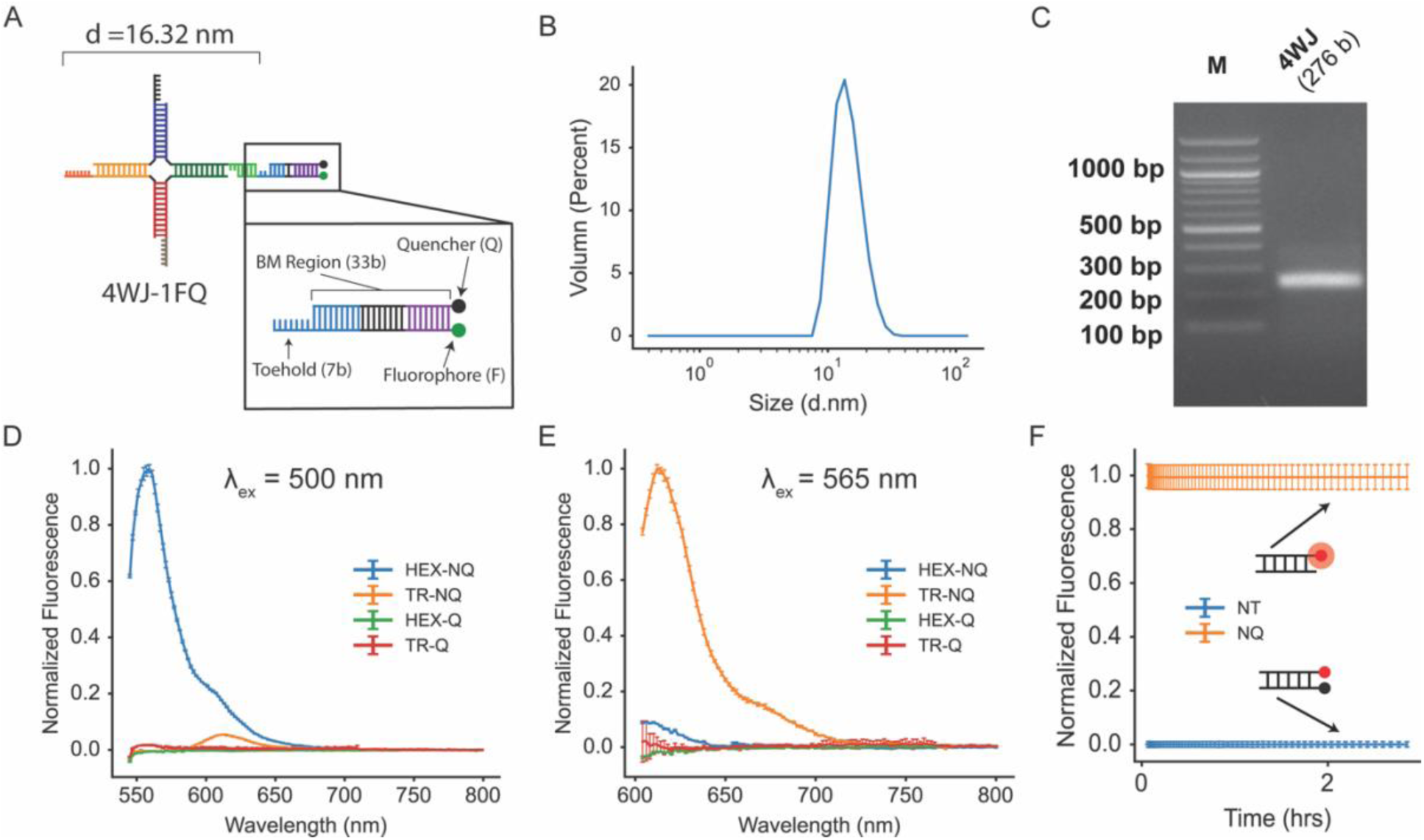
4-way junction (4WJ) schematic and characterization. (A) Schematic of the 4WJ with one fluorophore/quencher pair. (B) A representative volume percent curve from the DLS measurement of 4WJ. (C) Agarose gel image of the 4WJ. (D, E) Normalized fluorescent measurement of the quenched and unquenched system using HEX and TR excitation wavelength respectively. (F) Normalized fluorescent measurement for stability study.

### Trigger Design Screening

To further refine our previous strand displacement system design and offer a better control of the release kinetic, we chose to examine how the trigger’s sequence design could influence the displacement behaviour of the system. There are two main sequence design factors that can potentially use to influence the reaction kinetics, namely the mismatch type and mismatch location (32, 39). By incorporating mismatched base at varying location in the trigger strand, we aim to control the displacement properties of the strand displacement reaction system and allow us to further increase the versatility of our system and reduce the need for redesign. The mismatch type used in the study are presented in Table 1 and the mismatch locations are defined as 5’ to 3’ position of the mismatch appeared on the trigger strand. All the trigger strands used in the study can be found in **SI**

#### Evaluate and model the effect of various toehold region length and dangling bases on displacement quantity and rate

To compare our experimental results with existing model and data in the literature (28), we designed five triggers with varying toehold length (3-7 bases) and tested at 1:1 ratio (trigger:4WJ). The displacement behaviours for various triggers were fitted with both one-phase association and second order kinetic models. Two different types of models were used since we observed that the second order kinetic model deviated from the experimental result as we introduced mismatched bases into the system. A potential cause of the deviation is that the data collected is only based on the final product after complete displacement and since we introduced mismatched bases into the system, there are more likely intermediate complexes present that are not accounted for when solving the initial value problem on the system of equation. As expected, the increase in toehold length increased the overall reaction rate in both models, which is consistent to previous works and it is likely due to the higher affinity from the longer toehold (21, 23, 28–30, 32). Additionally, it has been shown that strand displacement reaction rates *in vitro* tend to plateau when the number of toehold bases reaches near eight nucleotides and our experimental fit results using the two models follow the same trend (Figure 2A, 2C). Furthermore, the rate obtained from the second order reaction model matched well with the second order reaction result from Zhang and Winfree (28). Welch one-way ANOVA revealed significant effect of the number of toehold bases on the rate constant for both second-order reaction model [F (4, 20.76) = 1995.4, p<0.001, η^2^ = 0.990] and one-phase association model [F (4, 25.59) = 1119.51, p<0.001, η ^2^ = 0.984]. Additionally, the relationship between the toehold bases and final displacement quantity also exhibits similar trend with the reaction rate, such that increase in toehold bases increase the final displacement quantity and plateaus as the toehold bases approaches seven toehold bases (Figure 2B). Indeed, Welch one-way ANOVA revealed that the number of toehold bases has a significant effect on the final displacement quantity based on one-phase association model [F (4, 23.68) = 892.163, p<0.001, η^2^ = 0.922]. The large η^2^ for all three parameters indicates that the number of toehold bases has a strong effect on those parameters. Furthermore, Games-Howell post hoc test was carried out for all three parameters and shown in Figure 2.

**Figure 2:**
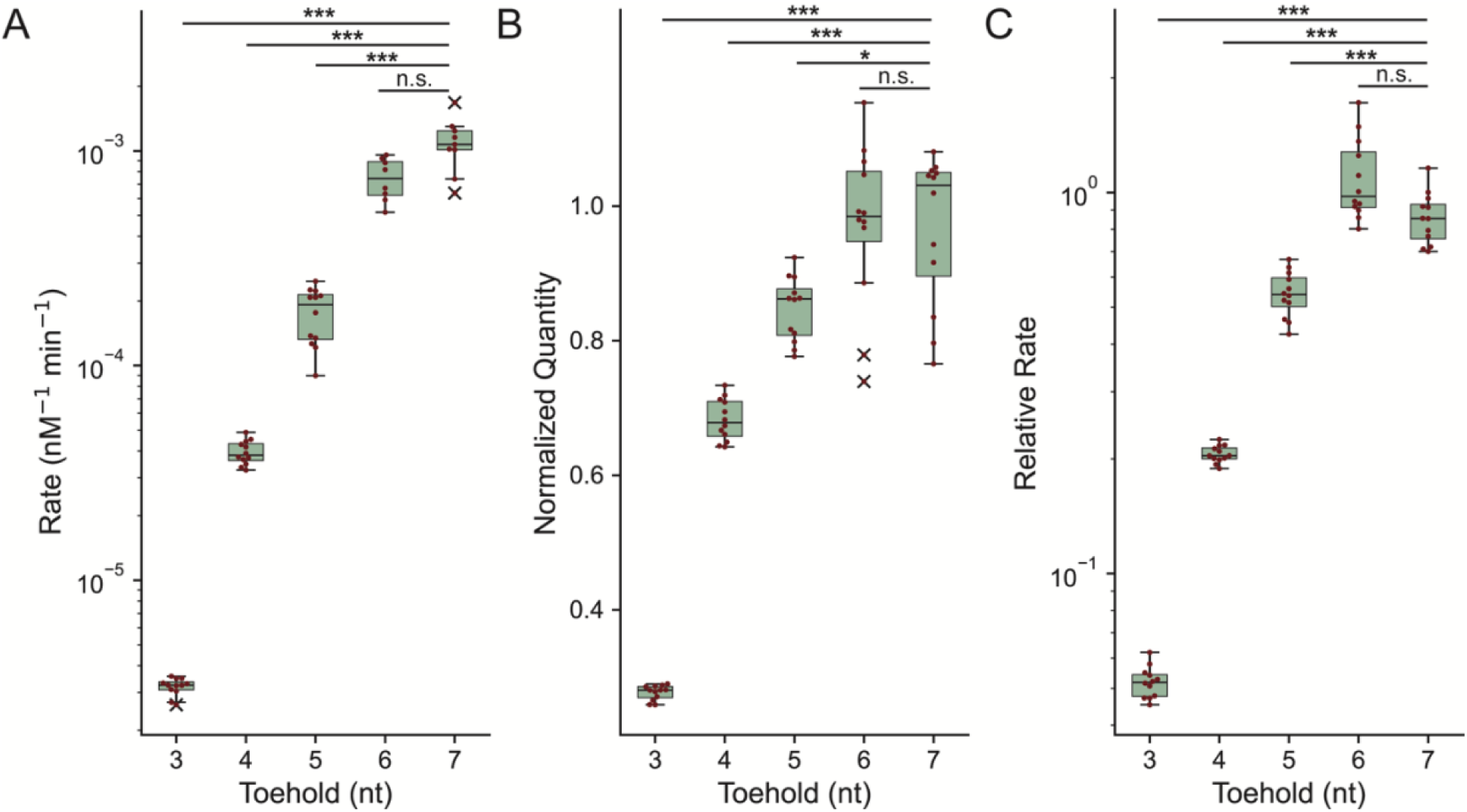
Boxplots of the fitted rates and quantity of triggers with varying toehold length using both second order reaction model and one-phase association model (ratio trigger:4WJ = 1:1). (A) The fitted reaction rate constant from second order reaction model. (B) The fitted final displacement quantity from the one-phase association model normalized against the displacement quantity of perfectly matched trigger. (C) The relative rate of the fitted reaction rate from the one-phase association model against perfectly matched trigger. (Welch one-way ANOVA with Games-Howell post-hoc analysis, n.s. = no significance, *=p<0.05, **=p<0.01, ***=p<0.001)

#### Evaluate and model the effect of mismatch conditions on displacement quantity and rate

After demonstrating that our system behaves similarly as the established system in literature (28), we incorporated various mismatch type at different location in effort to control the displacement properties of our system and modulate the release rate and quantity as suggested by previous work on strand displacement (31, 32). The mismatch types used in this study are presented in Table 1. We chose to include the mismatch base(s) on the trigger strand to avoid modification of the target strand, which facilitate comparison and would simplify future use of this system and reduce the cost associated with modification of the system. The mismatch location is defined as the location of mismatch base on the trigger strand with the base #1 being located on the 5’ end. By incorporating different types and number of mismatches at various locations, we expected to be able to slow down the displacement rate and lower the displacement quantity, which would allow us to define a set of rules and tune the exact displacement properties. For strand displacement modelling with mismatch presented, Seidel et. al. has utilized second order kinetic model with 1D Markov chain simulation and compared with the experimental data (32). To understand how the displacement behaviours are impacted by mismatches, the displacement behaviours for triggers with various mismatch conditions were modelled with one-phase association and second order kinetic model using experimental data. Both methods performed equally well and yield results that are relatively similar with the experimental final release quantity is comparable to the theoretical maximum release quantity (500 nM). However, as the experimental final release quantity deviates from the theoretical maximum released quantity, the one-phase association outperformed the kinetic models. This discrepancy is likely due to the kinetic model that is oversimplified with no intermediates present. Indeed, it has been shown that with the introduction of mismatched bases, the reaction kinetic for the system is likely to shift towards a second order reaction with a reversible intermediate step and an irreversible product step instead of a simple irreversible second order reaction (32). This observation was based on the thermodynamic simulation that was able to predict a specific pulse generation circuit that corresponds to existing experimental data. However, it is challenging to directly monitor both the presence of intermediate and displaced strands experimentally via fluorescent time point system only. Due to the difficulty to monitor both intermediate and displaced strands simultaneously in experiments, we limited our kinetic model fitting to a single irreversible release step.

One of the design parameters tested in the study was the type of mismatch presented in the trigger strand. The type of mismatched is defined as the base pair of the mismatched base and the corresponding base on the hybridizing strand (table 1). Additionally, the order of the pair matters in our definition, such that G:T and T:G are considered different mismatch types. Among the 12 different mismatch types outlined in Table 1, we only tested the first nine types due to the limitation in the sequence of the strands used in this study. Future study would include a larger sequence pool for both the trigger and target strands and examine how it impacts the release properties. The initial analyses were conducted on mismatch groups (Table 1), which is based on what the original base was, i.e., mismatch type 1-3 is the same group since the mismatched bases replaced the same initial base presented. All three mismatch groups were tested at the 5’ end near the start of branch migration region on the trigger strand (base 17 for group 1, base 18 for group 2, and base 15 for group 3). Trigger design schematic can be found in the SI (**Figure S2**). Among the three mismatch groups tested, both the plateau [F (2, 24.63) = 9.95, p<0.001, η ^2^ = 0.320 for group 1; F (2, 23.69) = 142.37, p<0.001, η ^2^ = 0.663 for group 2; F (2, 22.12) = 4.76, p<0.05, η ^2^ = 0.276 for group 3] and rate [F (2, 25.78) = 356.12, p<0.001, η ^2^ = 0.937 for group 1; F (2, 20.70) = 611.02, p<0.001, η ^2^ = 0.981 for group 2, F (2, 23.70) = 757.18, p<0.001, η ^2^ = 0.977 for group 3] differ significantly for the first three groups (type 1-9). Along with the Welch ANOVA analysis, Games-Howell post-hoc analysis was carried out for the parameters that are significant and the results along with both plateau and rate for all three groups and individual data points are presented in Figure 3. Based on the statistical result, the effect of mismatch type is more prominent on the rate compared to the plateau in all three mismatch groups. Moreover, the structural free energy and the theoretical concentration for each trigger-base pair were calculated using NUPACK as described in the method section (36). The calculated structural free energy among the nine trigger conditions differs by ∼±1.5 kcal/mol. Interestingly, the calculated structural free energy does not correlate with the result obtained in this study as the lower free energy should indicates the conformation is more favourable compared to the other conditions like the intermediate complex. This deviation might be caused by the reaction condition, which is not fully accounted for in the NUPACK simulation. However, the overall trend of the displacement behaviour for the mismatch type correlates well with the base pair stability reported in literature (39, 40). As indicated in the study by SantaLucia and Hicks, outside of the Watson-Crick pairs, guanine is the most tolerable mismatch base and cytosine is the least tolerable mismatch base with both adenine and thymine having a relatively similar effect between guanine and cytosine (39). This indicates that not only the presence of mismatch can impact the release properties but also which nucleotide is the mismatch can impact it. This indicates that it is indeed possilbe to tune the release profile of a release system by altering one single nucleotide in the system, which can vastly expand the potential design space for the strand displacement system.

**Figure 3:**
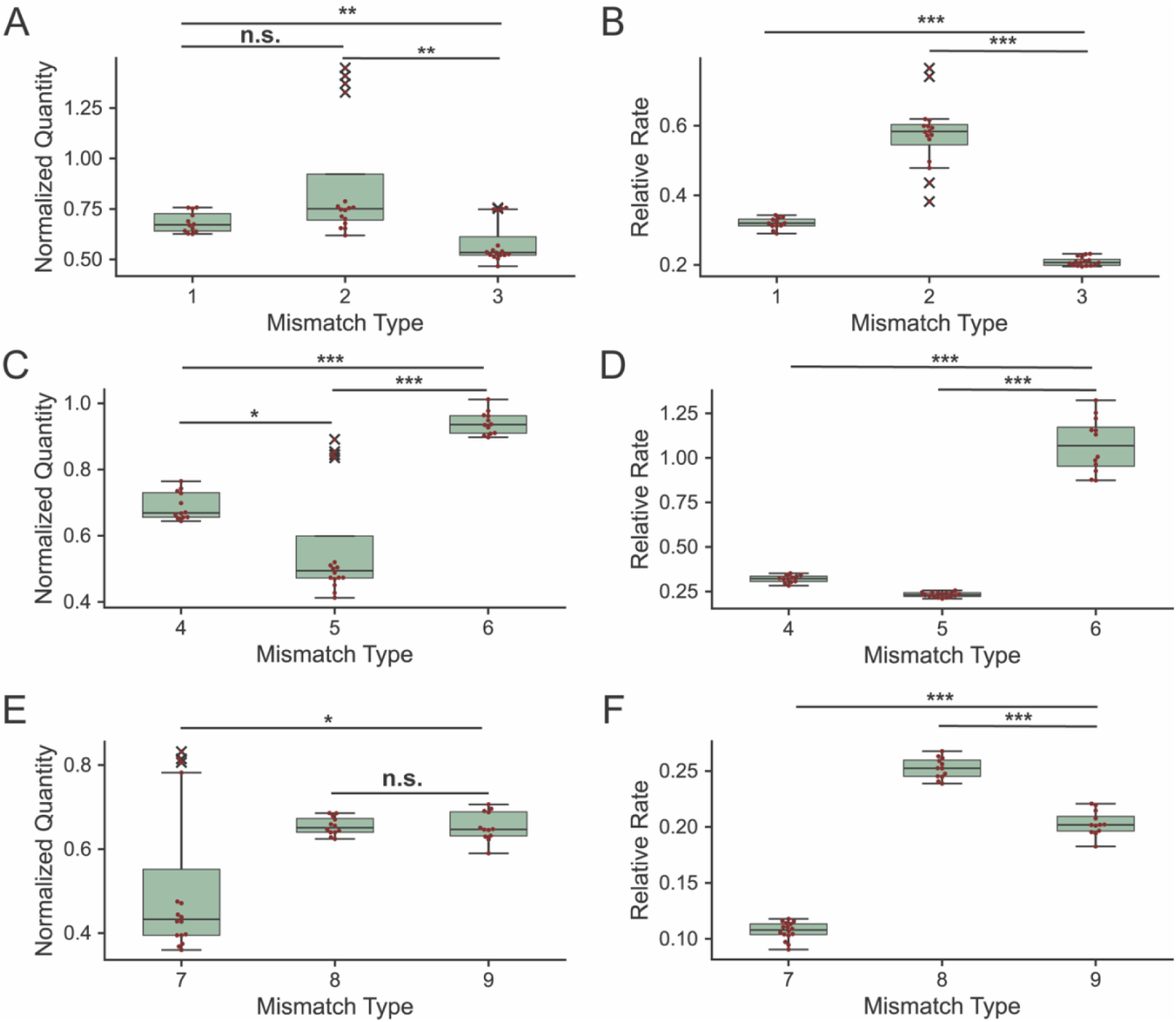
Boxplots of the fitted rates and quantity of triggers of the three mismatch groups using one-phase association model. The mismatch type number corresponds to the number shown in **Table 1**. (A, B) Normalized displacement quantity and relative rate for mismatch group 1 at location 17. (C, D) Normalized displacement quantity and relative rate for mismatch group 2 at location 18. (E, F) Normalized displacement quantity and relative rate for mismatch group 3 at location 15. Displacement quantity was normalized against the displacement quantity of perfectly matched trigger. The relative displacement was relative to the displacement rate of perfectly matched trigger. (Welch one-way ANOVA with Games-Howell post-hoc analysis, n.s. = no significance, *=p<0.05, **=p<0.01, ***=p<0.001)

Furthermore, we examined the effect of mismatch location on both displacement quantity and rate by using the same mismatch type along the trigger strand. Specifically, we used the type 1 mismatch (T:T) and found that the mismatch location has significant effect on both the plateau [F (5, 34.34) = 92.76, p<0.001, η^2^ = 0.475] and rate [F (5, 31.94) = 333.39, p<0.001, η^2^ = 0.966] for the system, which indicate that the mismatch type is not the only factor influencing the displacement behaviour (Figure 4A, B). The decreasing impact of mismatch on rate as mismatch location shift towards the end of branch migration region is like what was observed by Irmisch et. al. when they introduced mismatch at varying location on the trigger strand (32). This is probably due to the higher likelihood of the incumbent strand to self-dissociate from the target strand since the incumbent strand would have decreasing bounded base pair if the mismatch location on the trigger strand is located towards the end of the branch migration region. However, the lack of difference in the final displacement quantity was unexpected as one would suspect that the increasing change of incumbent self-dissociation would also result in a higher final displacement quantity. The change was only most prominent when the mismatch location was as position 33, which is two bases away from the end of the sequence. While it was unexpected, it does correlate well with the free energy calculated by NUPACK. The calculated free energy for the first five location variation of Type 1 mismatch was all ∼33 kcal/mol compared to the last variation, which is -35.68 kcal/mol. As the free energy would dictate the final equilibrium concentration of the system, the lack of different in the first five location variation of Type 1 mismatch should be expected based on the free energy calculation. To further examine how mismatch type and location influence the displacement behaviour, mismatch group 1 and group 3 were used. Each member of the mismatch group was used in two different locations, one near the start of the branch migration region and another near the end of it. For the mismatch group 1, they were inserted into base position 17 and 29. For mismatch group 3, they were inserted into base position 15 and 27. Two-sample T-test with unequal variance assumption was carried out on individual mismatch type pairs and Hedge’s ‘g’ was calculated. Interestingly, the plateau was only significantly different for three pairs of mismatch types (mismatch type 6, 7, and 9) compared to the rate, which all mismatch pairs were significant (Figure 4C, D, E, F). This result correlates to the previous result that both mismatch type and location has a weaker effect on the plateau compared to the rate. The calculated free energy from NUPACK for the Trigger 1 system can be found in the SI (**Table S3**).

**Figure 4:**
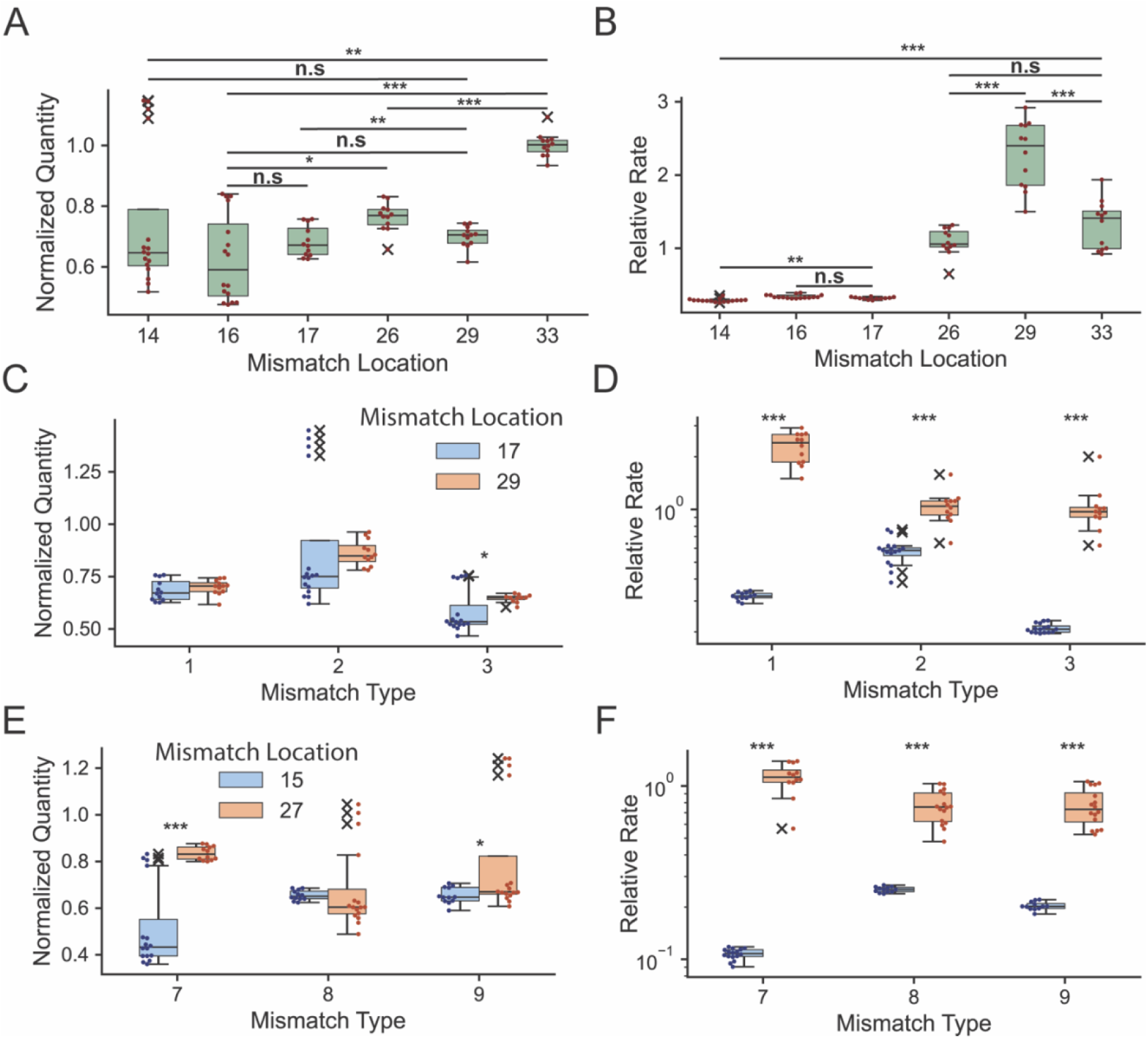
Boxplots of the fitted final displacement quantity and rate for varying trigger designed based on mismatch location. (A, B) Type 1 mismatch (T:T) at varying base location using one-phase association model. (Welch one-way ANOVA with Games-Howell post-hoc analysis, n.s. = no significance, *=p<0.05, **=p<0.01, ***=p<0.001). (C, D) Mismatch group 1 (type 1, 2, 3) at location 17 and 29 (two sample T-test with unequal variance, *=p<0.05, **=p<0.01, ***=p<0.001). (E, F) Mismatch group 3 (type 7, 8, 9) at location 15 and 27 (two sample T-test with unequal variance, *=p<0.05, **=p<0.01, ***=p<0.001). Displacement quantity was normalized against the displacement quantity of perfectly matched trigger. The relative displacement was relative to the displacement rate of perfectly matched trigger.

An important assumption that we made in our definition for the mismatch type is that the order of the mismatch matters and the displacement behaviour for G:T mismatch type would significantly differ from T:G mismatch type. In effort to examine this assumption, mismatch type 2 (G:T) and mismatch type 8 (T:G) at various base location were examined together by both base location and mismatch type. There is a total of three different locations (base 17, 21, and 29) for mismatch type 2 and four different locations (base 13, 15, 27, and 31) for mismatch type 8. For the base location, we assume that the two types of mismatches are the same and the only factor that would influence the displacement behaviour is the position of the mismatch base on the trigger strand. Welch one-way ANOVA indicated that the mismatch location has a significant effect on both plateau [F (6, 38.76) = 132.56, p<0.001, η ^2^ = 0.305] and rate [F (6, 36.51) = 358.93, p<0.001, η ^2^ = 0.926] obtained from the one-phase association model. For the mismatch type analysis, the values from various locations of same mismatch type are pooled together and a two-sample T-test with unequal variance along with Hedge’s g was carried out. The two-sample T-test indicates that both plateau [two-sample t(67.86) = 5.93, p<0.001, g = 1.28] and rate [two-sample t(80.49) = 4.06, p<0.001, g = 0.743] are significantly different between the two mismatch types even across varying mismatch location. Therefore, it is reasonable to assume that the order of mismatch does influence the displacement behaviour of the system and should be considered as two different types of mismatches. The boxplot showing both mismatch location and type analysis can be found in SI (**Figure S3**).

#### Evaluate the effect of two-mismatch trigger design on displacement quantity and rate

Addition to the one mismatch trigger design, we also examined trigger design with two mismatches presented. For the two-mismatch trigger design, the order of the mismatch presented was examined using consecutive mismatch that are both in group 1 mismatch defined in Table 1. It is found that the type of mismatch at the second location has a significant impact on both released quantity [F (2, 21.35) = 821.55, p<0.001, η ^2^ = 0.981; F (2, 20.57) = 696.61, p<0.001, η ^2^ = 0.985; F (2, 19.75) = 829.12, p<0.001, η ^2^ = 0.973] and rate [F (2, 20.87) = 647.96, p<0.001, η ^2^ = 0.970; F (2, 21.37) = 1430.17, p<0.001, η ^2^ = 0.990; F (2, 21.90) = 16.58, p<0.001, η ^2^ = 0.514] except for one pair (Figure 5A, B). Furthermore, the change in the release quantity and rate are similar to the base pairing stability study from SantaLucia and Hicks (39). In addition to the combination of mismatch types, the effect of second mismatch location was also examine by fixing the first mismatch (type 8) at location 13 and the second mismatch (type 1) at varying position (Figure 5C, D). Both release quantity [F (2, 19.29) = 73.28, p<0.001, η ^2^ = 0.857] and rate [F (2, 20.66) = 2512.96, p<0.001, η ^2^ = 0.991] were significantly impacted by where the second mismatch location was located. The rate obtained from the one-phase association model follows the similar trend as the two-mismatch system simulation conducted by Irmisch et. al. assuming the closed configuration for the displacement reaction (32). Interestingly, the release quantity follows a different a pattern that shows the consecutive mismatch result in a higher final release quantity and the trigger with shorter mismatch spacing has a lower release quantity compared to the trigger with longer mismatch spacing. An important caveat for the two-mismatch system is that some of the trigger system did not fully reach plateau at the end of the experimental time; therefore, it would likely need to be study further with a longer experimental time to allow it to fully reach the plateau region. However, it demonstrates the possibility of a slow-release system with TMSDR using pure DNA components without any chemical or biological alteration of the system.

**Figure 5:**
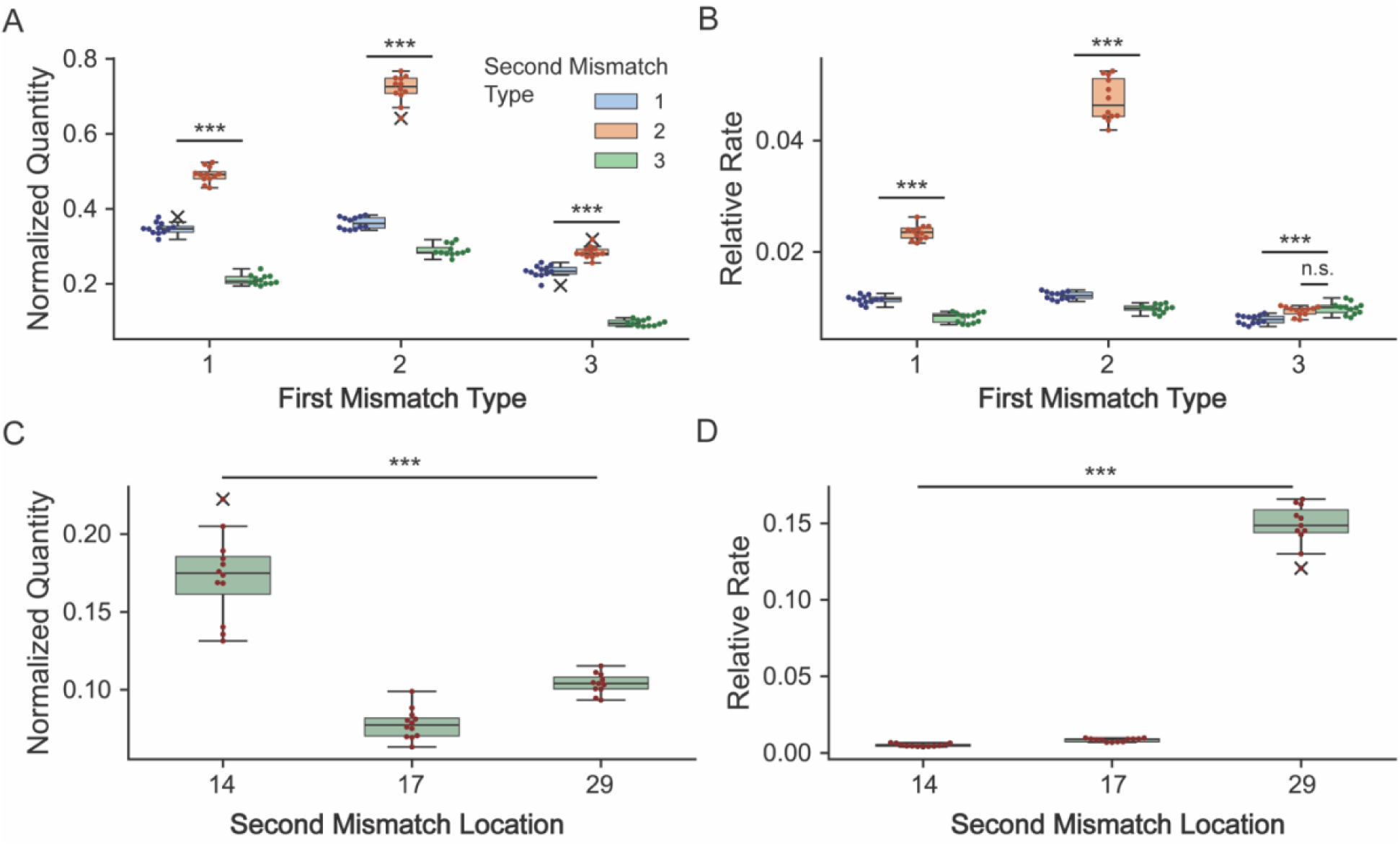
Boxplots of the fitted final displacement quantity and rate for two-mismatch trigger design. (A, B) Two-mismatch trigger design with consecutive mismatch using one-phase association model. (C, D) Two-mismatch trigger design with type 8 mismatch at location 13 and type 1 mismatch at varying location. (Welch one-way ANOVA with Games-Howell post-hoc analysis, n.s. = no significance, ***=p<0.001).

#### Evaluate and model the effect of trigger concentration on displacement quantity and rate

To assess the influence of the trigger concentration on hybridization quantity, the trigger concentration was varied from 0.25x to 2x trigger to target ratio with 0.25x increments and from 1x to 5x trigger to target ratio with 1x increments. This was carried out with three different triggers: Trigger 1 (**T1**), the perfectly matched trigger with 5 dangle bases and 7 toehold bases, A1 (Type 1 mismatch at position 17), and A2 (Type 1 mismatch at position 17 and Type 6 mismatch at position 18). The rate obtained using a one phase association model with these different ratios shows a concentration dependent behaviour (Figure 6A, B, C); however, it is important to note that the concentration dependent behaviour varied significantly between the different types of triggers tested. For instance, T1 exhibited a decrease in the overall rate as its concentration approaches 1x molar ratio against the target and remained relatively steady as the concentration increases. The trigger A1 and A2, however, exhibited an opposite behaviour compared to T1. Both rate from A1 and A2 were relatively steady until the trigger concentration reached ∼1.5x molar ratio against the target and the rate increased with increasing trigger concentration. This shift in behaviour was interesting and unexpected, which will be examine further in a future study. To try to understand this results, we examine the raw data and we observed that at lower concentration of T1, the release curves always exhibit a steep increase at the initial stage (from 0 to 10 minutes). For the one-phase association model, the rate is calculated based on the time to reach the plateau, a combination of steeper initial release and a lower plateau value could contribute towards the higher rate of T1 at a lower concentration. In contrast with T1, we observed a relatively less steep initial stage for A1 and a shallow initial stage for A2. The steepness of the initial stage for both A1 and A2 does seem to increase as the concentration increases, which could lead the increase in rate as observed. This shift of behaviour in the rate observed could potentially cause by the experimental conditions. On the other hand, the hybridization quantity showed a consistent concentration dependent behaviour up the point where the final hybridization quantity is near the theoretical max hybridization quantity (Figure 6D, E, F). This is most clearly shown in Figure 6D with the perfect trigger strand that reached the theoretical max hybridization quantity at 1x trigger to 4WJ ratio. The relationships between the trigger:4WJ ratio and hybridization quantity were fitted using a simplified one-phase association model without lag [equation (2)]. The fitted parameter values for each trigger can be found in **SI**. This concentration-dependent behaviour of total release quantity is expected; however, it is harder to predict the relationship between trigger:4WJ ratio and the final release quantity once mismatched base(s) are included in the system without some empirical data since an unknown quantity of the intermediate complex would be present in the system. A caveat for the final release quantity of Trigger A2 is that based on the release curve generated for individual trials, it did not fully reach the plateau within the duration of the study. Therefore, a longer study duration would be carried out for the future study for trigger designs that includes more than one mismatched base.

**Figure 6:**
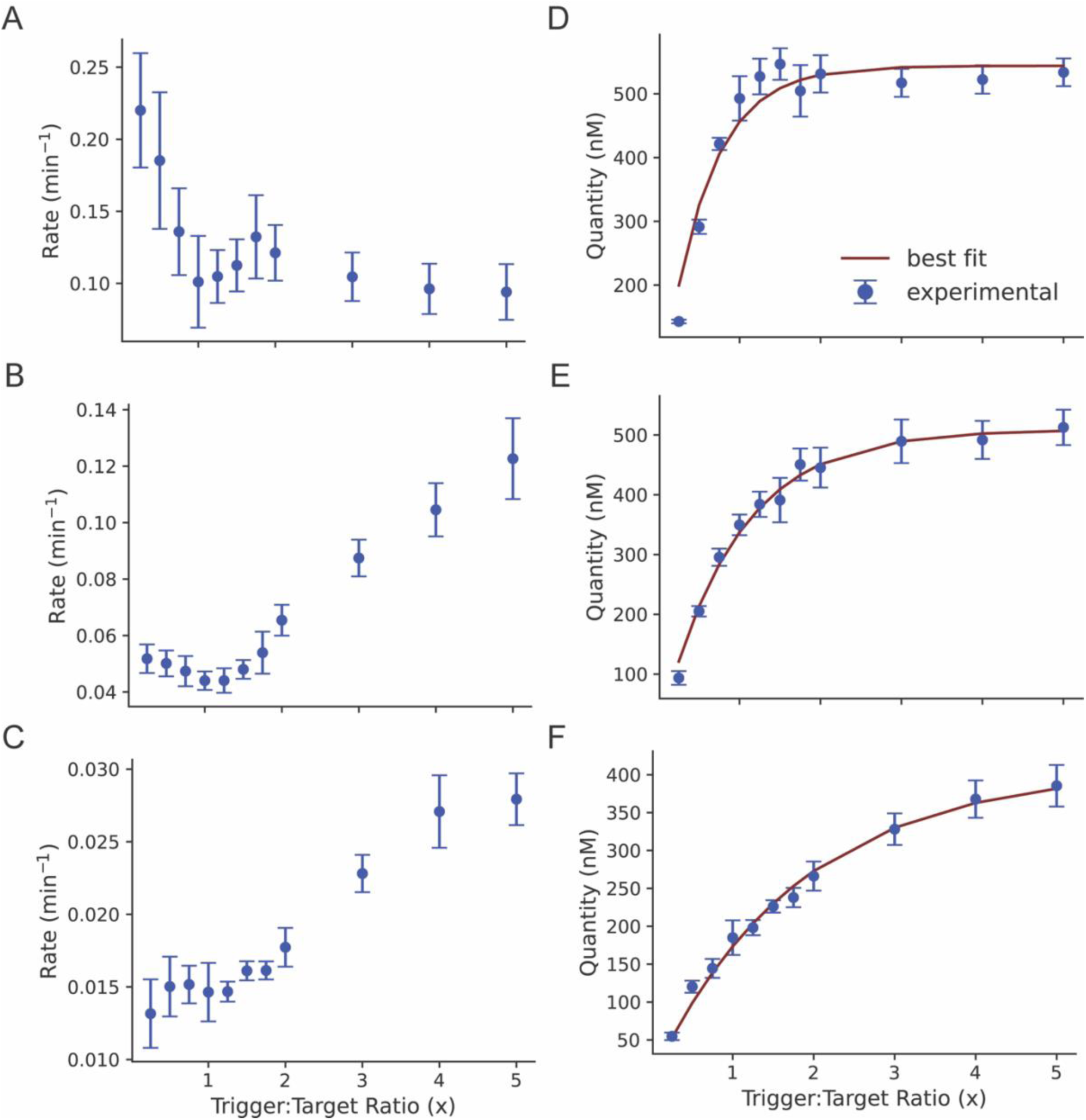
The hybridization parameters determined by one-phase association model at varying trigger:4WJ concentration ratio. Each point is presented as mean *±* SD. (A, B, C) The rate values determined at various trigger-to-target ratios. (D, E, F) The plateau values determined at various trigger:4WJ concentration ratios with the best fitted line using the one-phase association model. (A, D) The perfectly matched trigger. (B, E) Trigger A1 with one mismatched base at location 17 (T:T). (C, F) Trigger A2 with two mismatched bases at location 17 and 18 (T:T and G:A).

### Toward Predicting the Release Behaviour of the System

#### Predicting the displacement behaviour of a similar trigger design

After the initial screening of the designed trigger sequence, we aim to utilize the displacement behaviour information gathered in the previous section to develop a model to predict the displacement behaviour of similarly designed trigger. The displacement model is developed with previously defined parameters (mismatch type and location) using linear regression. To examine the accuracy of the model, a series of triggers were designed with the defined parameters (mismatch type and location) and examined in the same method as the screening stage and the box plot of each condition can be found in **SI**. The design parameters were feed to the model to generate the predicted displacement properties and compared with the experimentally generated displacement properties.

The same series of triggers was also simulated in NUPACK to predict the theoretical displacement quantity in a tube at the specific testing condition. Most of the predicted results from NUPACK were overestimating compared to the experimental results while the regression model has a mix of over- and underestimation (Figure 7A). Since NUPACK based its calculation on the calculated theoretical free energy for each duplex, it is unsurprising that the concentration calculations for all test strands have similar values. However, it is surprising that the decreasing in free energy caused by the introduction of mismatch (∼17%) did not influence the theoretical concentration calculated in any significant manner (∼2% decrease). The calculated free energy from NUPACK can be found in the SI (**Table S7**). It is possible that the inclusion of the fluorophore and quencher in the experimental system could cause the deviation from the NUPACK with experimental data collected as previously described in literature (41, 42). Since our model is based on the experimental data from our system, the effect of fluorophore and quencher should be less pronounced; however, the type of fluorophores and quenchers does also impact the stability of the system (41, 42), thus, some of the deviation observed in the prediction might be partially due to the change in fluorophore and quencher used in the system for the different trigger strands tested. Overall, the displacement quantity prediction has an average of -6% difference between the prediction and the experimental result compared to the 20% from NUPACK. Additionally, the NUPACK simulation has generally overestimated the final release quantity (17/18 samples overestimated) compared to the regression model (5/18 sample overestimated).

**Figure 7:**
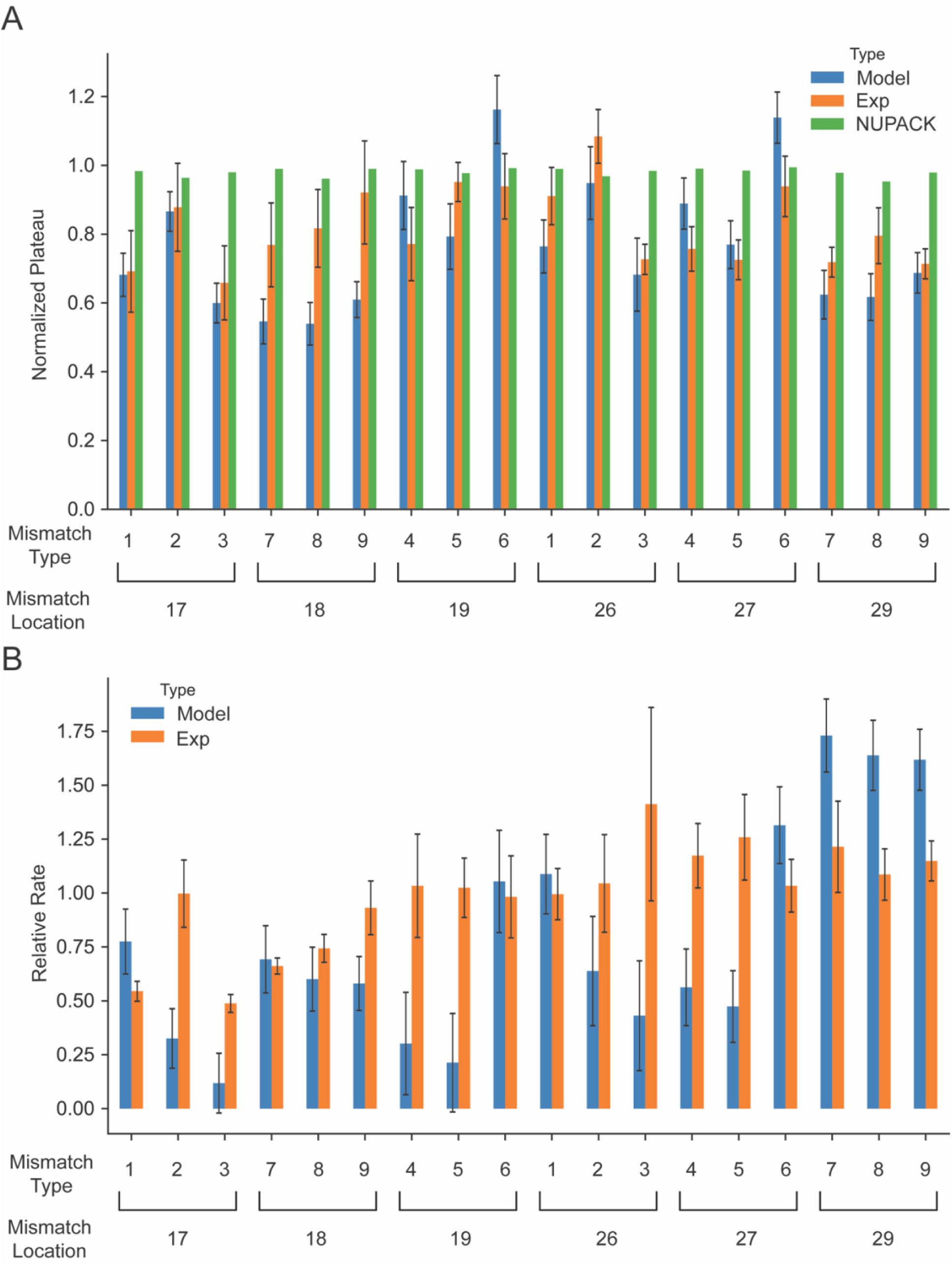
The experimental result and the prediction result from linear regression with dummy encoding and NUPACK simulation. (A) Normalized plateau from experimental data, linear regression model, and NUPACK simulation. (B) Relative rate from experimental data and linear regression model. The experimental data was represented as mean ± std. The linear regression model result was represented as mean prediction ± 95% confidence interval.

Furthermore, root mean square error (**RMSE**) for the two models were calculated (RMSE_reg_=0.162; RMSE_nupack_=0.190), which indicates the linear regression with dummy encoding outperforms the NUPACK simulation slightly. However, only ∼33% of the experimental mean fall within the 95% confident interval of the linear regression model. Overall, this indicates that NUPACK can be a good starting point for trigger design, but the regression model would be likely be more useful for designing triggers that are similar in sequence with mismatch bases included.

In addition to the displacement quantity, displacement rate was also modelled using the same method as the displacement quantity. However, the displacement rate prediction did not perform as well compared to the displacement quantity prediction (Figure 7B). Similarly to the displacement quantity prediction, the displacement rate prediction is a mixture of over- and under-estimation with ∼44% of the predictions are overestimation and an average of -19% difference between the prediction and the experimental result and the RMES of 0.526. The poorer performance of the rate model could be due to both overall quality and quantity of the model data, which could be potentially alleviated by conducting more replicates of existing condition and introducing more conditions into the sampling data to further improves the model. This would likely lead to improvement in both release quantity and rate model.

### Utilizing the Mismatch Trigger Design for Multiplex Release System

In addition to examine and model the mismatch trigger design, we also used to examine the possibility of using the mismatch trigger design to modulate multiplex release system. It was shown in our previous work that it is possible to release two separate cargos using the 4WJ DNA motif with the perfectly matched triggers (19). However, using the mismatched trigger to generate a different release pattern for multiplexed release was not shown. The trigger solution was prepared with two trigger strands, one for HEX fluorophore and one for TR fluorophore, at 1:1 trigger:4WJ ratio. Two different mismatch triggers were selected for this study, A1 and A2. Trigger A1 is like T3 but with type 1 mismatch at location 17. Trigger A2 is like T1 but with two mismatches, type 1 mismatch at location 16 and type 6 mismatch at location 17. As shown in Figure 8A, B, the mixture of T1/T3 behaves similarly to the mixtures when T1 or T3 were present individually for both the quantity and rate. Additionally, the presence of mismatch triggers did influence the release quantity and rate without impacting the release profile of the other cargos. Furthermore, the release curve for both HEX and TR can be seen in Figure 8C, D, E. This indicates that the use of mismatched trigger is not limited to only singular use, but by using a trigger mixture, such as regular trigger with mismatched trigger, it can be used to finely tune the behaviour of the multiplexed system.

**Figure 8:**
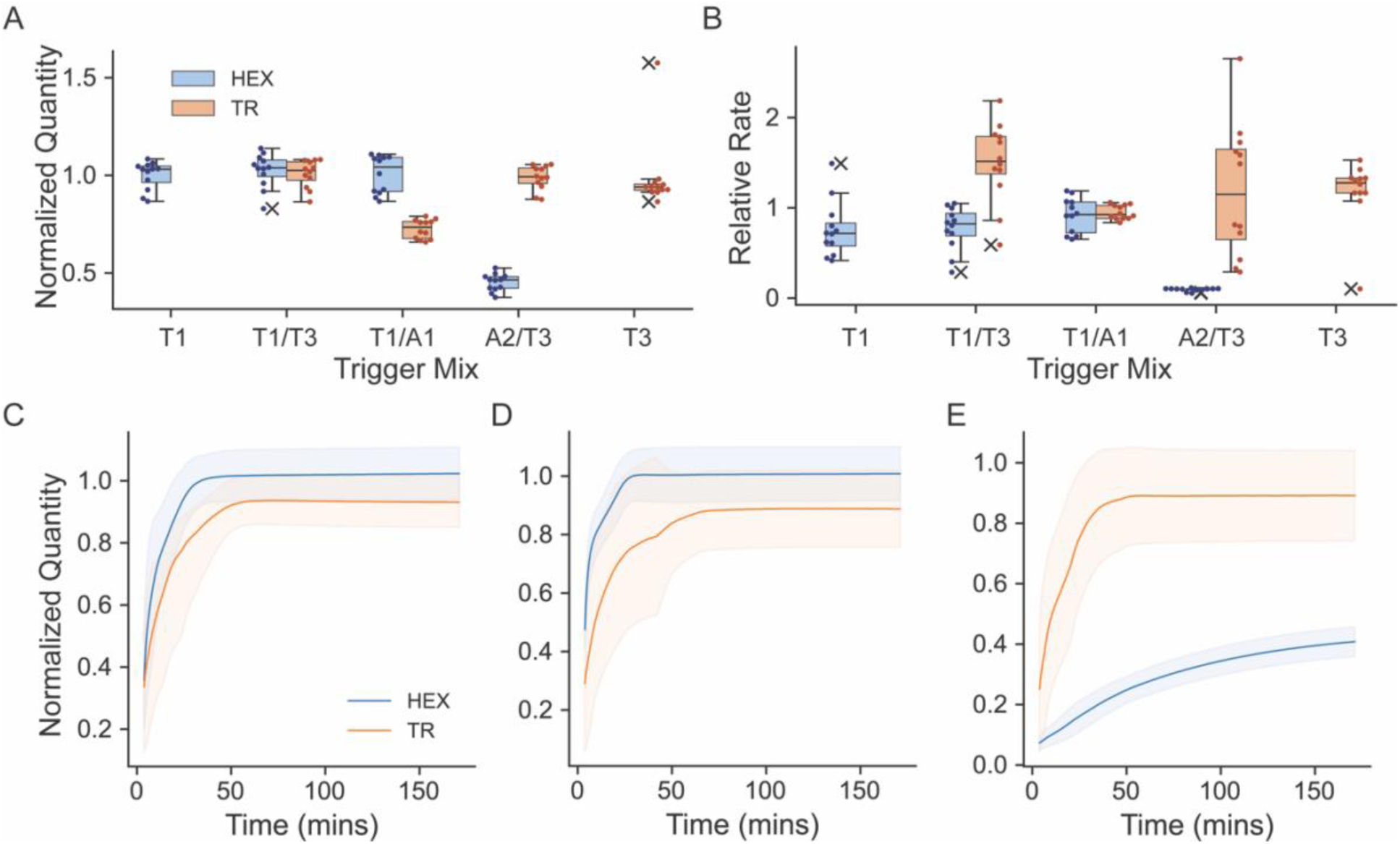
Multiplexed release with mismatched trigger system. (A, B) The release properties of each combination of trigger mixture for both HEX and TR fluorophore. (C, D, E) The release curve for T1/T3, T1/A1, A2/T3 mixture, respectively. The curve was represented with mean ± std.

## Conclusion

The design rules for DNA toehold-mediated strand displacement reaction (**TMSDR**) utilizing the presence of mismatched base(s) in the trigger was investigated and modelled using linear regression with dummy encoding. It is shown that the location, type, and quantity of the mismatch can significantly impact both the release rate and quantity of the system. Using the linear regression with dummy encoding, we were able to predict some of the release properties of a similarly designed trigger strand. However, more work would be required to include a wider sequence pool and larger data sets to improve the model prediction for trigger design. In addition to the investigating and modelling how the presence of mismatch influence on the TMSDR, we also investigated the concentration-dependent behaviour of both perfectly matched trigger and mismatched triggers. We found that both the release quantity and rate were dependent on the concentration; however, the concentration-dependency of the release rate were also trigger-dependent. To better understand the implications of this later observation, it would require further study with a larger sequence pool at varying concentration to further understand the relationship between concentration, trigger design, and the release rate.

In addition to investigating and modelling the mismatch trigger system, a DNA motif (**4WJ**) with capability of multiplexed release function were fabricated and examined using fluorophore/quencher pairs. The multiplexed release function was further expanded from the previous study by incorporation of the trigger mixture with both regular and mismatch triggers. The incorporation of the mismatch trigger in the trigger mix was able to modulate the release of specific cargo without impacting on the release of other cargos. This further expand the capacity of the multiplex system with the ability to finetune the release property without complete redesign of the system or addition of additional chemical or biological components.

Supplementary Data are available at NAR online.

## AUTHOR CONTRIBUTIONS

Remi Veneziano: Conceptualization, Experimental Design, Data Analysis, Visualization, Writing—review & editing. Chih-Hsiang Hu: Conceptualization, Experimental Design, Data Collection and Analysis, Visualization, Writing—original draft, review & edit.

## ACKNOWLEDGEMENTS

C-H. H gratefully acknowledge the Department of Bioengineering at George Mason University for its support.

