## Supplemental Information for "Designer DNA Strand Displacement Reaction toward Controlled Release of Cargos"

**MANUSCRIPT TITLE**

The title must be clearly intelligible to a non-specialist. The use of jargon and non-standard abbreviations in the title is not permitted

**AUTHORS**

Read the full [Authorship Guidelines](https://academic.oup.com/pages/authoring/journals/preparing_your_manuscript/ethics#Authorship).

Chih-Hsiang Hu^1^, Remi Veneziano^1,^*

^1^ Department, Institution, Town, State, Postcode, Country

* To whom correspondence should be addressed. Tel: [insert phone number including country code]; Fax: [insert fax number including country code]; Email: [insert email address]

**SUPPLEMENTARY DATA**

Table S1: DNA sequences (5’ to 3’) used for 4-way junction (**4WJ**) and perfectly matched trigger.

| Name | Sequence |
| --- | --- |
| 4WJ_1 | CTG ACC GAG CGT GGC TAC AGC TTC TTT GGT TCA CCG CTT GCC TTC TGC TCT CGC TGT GGC ACC TGC ACG |
| 4WJ_2 | CCT GCT TGC TGA TCC ACA TCT GCT TTT GAA GCT GTA GCC ACG CTC GGT CAG GCA TCG GGT ATC CAG TGG |
| 4WJ_3 | AGA GCA GAA GGC AAG CGG TGA ACC TTT CGT GAC TGC ACC TGA ATT GGC ACT TGC GCC ATC CCT TCA GAG |
| 4WJ_4 | AGT GCC AAT TCA GGT GCA GTC ACG TTT AGC AGA TGT GGA TCA GCA AGC AGG TAC GGT CGG CGC TGT ATG |
| Payload 1 HEX | /5HEX/ TG GGTGG TG AGA TG GATTG TG AGA TG TG AGA CAT ACA GCG CCG ACC GTA |
| Payload 1 Quench | CA CAATC CA TCT CA CCACC CA /3IABkFQ/ |
| Trigger 1 (T1) | CA TAACA CA TCT CA CAATC CA TCT CA CCACC CA |
| Payload 2 TR | /5TexRd-XN/ TG TTGGA TG AGA TG AGTTT TG AGA TG GG AGA CTC TGA AGG GAT GGC GCA |
| Payload 2 Quench | CA AAACT CA TCT CA TCCAA CA /3IABRQSp/ |
| Trigger 2 (T2) | CAC CAC CCA TCT CAA AAC TCA TCT CAT CCA ACA |

Table S2: DNA sequence (5’ to 3’) for the triggers used in toehold length study.

| Name | Sequence | Toehold length | ΔG (kcal/mol) |
| --- | --- | --- | --- |
| T1_tb3 | TCT CA CAATC CA TCT CA CCACC CA | 3 | -31.58372255 |
| T1_tb4 | A TCT CA CAATC CA TCT CA CCACC CA | 4 | -31.97088731 |
| T1_tb5 | CA TCT CA CAATC CA TCT CA CCACC CA | 5 | -32.98297171 |
| T1_tb6 | A CA TCT CA CAATC CA TCT CA CCACC CA | 6 | -33.94593103 |
| T1_tb7 | CA CA TCT CA CAATC CA TCT CA CCACC CA | 7 | -35.22305541 |

Table S3: DNA sequence (5’ to 3’) for 1 mismatched variation of Trigger 1 used in screening study. The location and type refer to the base location (5’ to 3’) and type of mismatch presented.

| Sequence | Location | Type | ΔG (kcal/mol) |
| --- | --- | --- | --- |
| CA TAACA CA TCT TA CAATC CA TCT CA CCACC CA | 13 | 8 | -31.670 |
| CA TAACA CA TCT CT CAATC CA TCT CA CCACC CA | 14 | 1 | -33.108 |
| CA TAACA CA TCT CA AAATC CA TCT CA CCACC CA | 15 | 7 | -32.650 |
| CA TAACA CA TCT CA TAATC CA TCT CA CCACC CA | 15 | 8 | -31.527 |
| CA TAACA CA TCT CA GAATC CA TCT CA CCACC CA | 15 | 9 | -33.172 |
| CA TAACA CA TCT CA CTATC CA TCT CA CCACC CA | 16 | 1 | -33.335 |
| CA TAACA CA TCT CA CATTC CA TCT CA CCACC CA | 17 | 1 | -33.100 |
| CA TAACA CA TCT CA CAGTC CA TCT CA CCACC CA | 17 | 2 | -32.525 |
| CA TAACA CA TCT CA CACTC CA TCT CA CCACC CA | 17 | 3 | -33.137 |
| CA TAACA CA TCT CA CAAAC CA TCT CA CCACC CA | 18 | 4 | -33.415 |
| CA TAACA CA TCT CA CAACC CA TCT CA CCACC CA | 18 | 5 | -32.755 |
| CA TAACA CA TCT CA CAAGC CA TCT CA CCACC CA | 18 | 6 | -34.424 |
| CA TAACA CA TCT CA CAATG CA TCT CA CCACC CA | 19 | 9 | -33.866 |
| CA TAACA CA TCT CA CAATC GA TCT CA CCACC CA | 20 | 9 | -32.739 |
| CA TAACA CA TCT CA CAATC CG TCT CA CCACC CA | 21 | 2 | -32.120 |
| CA TAACA CA TCT CA CAATC CA TCT GA CCACC CA | 25 | 9 | -32.792 |
| CA TAACA CA TCT CA CAATC CA TCT CT CCACC CA | 26 | 1 | -33.112 |
| CA TAACA CA TCT CA CAATC CA TCT CA ACACC CA | 27 | 7 | -33.485 |
| CA TAACA CA TCT CA CAATC CA TCT CA TCACC CA | 27 | 8 | -31.135 |
| CA TAACA CA TCT CA CAATC CA TCT CA GCACC CA | 27 | 9 | -34.258 |
| CA TAACA CA TCT CA CAATC CA TCT CA CAACC CA | 28 | 7 | -32.358 |
| CA TAACA CA TCT CA CAATC CA TCT CA CCTCC CA | 29 | 1 | -33.120 |
| CA TAACA CA TCT CA CAATC CA TCT CA CCGCC CA | 29 | 2 | -31.650 |
| CA TAACA CA TCT CA CAATC CA TCT CA CCCCC CA | 29 | 3 | -32.127 |
| CA TAACA CA TCT CA CAATC CA TCT CA CCAGC CA | 30 | 9 | -34.261 |
| CA TAACA CA TCT CA CAATC CA TCT CA CCACT CA | 31 | 8 | -32.446 |
| CA TAACA CA TCT CA CAATC CA TCT CA CCACC GA | 32 | 9 | -33.789 |
| CA TAACA CA TCT CA CAATC CA TCT CA CCACC CT | 33 | 1 | -35.681 |

Table S4: DNA sequence (5’ to 3’) for 2 mismatched variation of Trigger 1 used in screening study. The location and type refer to the base location (5’ to 3’) and type of mismatch presented.

| Sequence | Location | Type | ΔG (kcal/mol) |
| --- | --- | --- | --- |
| CA TAACA CA TCT CA CTTTC CA TCT CA CCACC CA | 16, 17 | 1, 1 | -31.585 |
| CA TAACA CA TCT CA CGGTC CA TCT CA CCACC CA | 16, 17 | 2, 2 | -30.605 |
| CA TAACA CA TCT CA CCCTC CA TCT CA CCACC CA | 16, 17 | 3, 3 | -31.185 |
| CA TAACA CA TCT CA CTCTC CA TCT CA CCACC CA | 16, 17 | 1, 3 | -31.385 |
| CA TAACA CA TCT CA CCTTC CA TCT CA CCACC CA | 16, 17 | 3, 1 | -31.386 |
| CA TAACA CA TCT CA CGTTC CA TCT CA CCACC CA | 16, 17 | 2, 1 | -31.188 |
| CA TAACA CA TCT CA CTGTC CA TCT CA CCACC CA | 16, 17 | 1, 2 | -30.987 |
| CA TAACA CA TCT CA CCGTC CA TCT CA CCACC CA | 16, 17 | 3, 2 | -30.788 |
| CA TAACA CA TCT CA CGCTC CA TCT CA CCACC CA | 16, 17 | 2, 3 | -30.992 |

Table S5: DNA sequence (5’ to 3’) for the triggers used in concentration study.

| Name | Sequence | Location | Type | ΔG (kcal/mol) |
| --- | --- | --- | --- | --- |
| T1 | CA TAACA CA TCT CA CAATC CA TCT CA CCACC CA | N/A | N/A |  |
| A1 | CA TAACA CA TCT CA CATTC CA TCT CA CCACC CA | 17 | 1 |  |
| A2 | CA TAACA CA TCT CA CATGC CA TCT CA CCACC CA | 17, 18 | 1, 6 |  |

Table S6: The fitted parameters for the hybridization quantity of concentration study. The parameters are reported with the value ± SE.

| Trigger | Plateau (nM) | Rate (unitless) |
| --- | --- | --- |
| T1 | 543.999 ± 16.189 | 1.828 ± 0.214 |
| A1 | 509.123 ± 9.417 | 1.086 ± 0.0579 |
| A2 | 406.967 ± 12.148 | 0.555 ± 0.0346 |

Table S7: DNA sequence (5’ to 3’) for 1 mismatched variation of Trigger 2 used in the prediction section. The location and type refer to the base location (5’ to 3’) and type of mismatch presented.

| Sequence | Location | Type | ΔG (kcal/mol) |
| --- | --- | --- | --- |
| CA CCACC CA TCT CA AATCT CA TCT CA TCCAA CA | 17 | 1 | -30.732 |
| CA CCACC CA TCT CA AAGCT CA TCT CA TCCAA CA | 17 | 2 | -29.939 |
| CA CCACC CA TCT CA AACCT CA TCT CA TCCAA CA | 17 | 3 | -30.366 |
| CA CCACC CA TCT CA AAATT CA TCT CA TCCAA CA | 18 | 8 | -29.649 |
| CA CCACC CA TCT CA AAAGT CA TCT CA TCCAA CA | 18 | 9 | -31.809 |
| CA CCACC CA TCT CA AAAAT CA TCT CA TCCAA CA | 18 | 7 | -31.440 |
| CA CCACC CA TCT CA AAACG CA TCT CA TCCAA CA | 19 | 6 | -31.849 |
| CA CCACC CA TCT CA AAACC CA TCT CA TCCAA CA | 19 | 5 | -30.245 |
| CA CCACC CA TCT CA AAACA CA TCT CA TCCAA CA | 19 | 4 | -31.127 |
| CA CCACC CA TCT CA AAACT CA TCT CT TCCAA CA | 26 | 1 | -31.368 |
| CA CCACC CA TCT CA AAACT CA TCT CG TCCAA CA | 26 | 2 | -30.077 |
| CA CCACC CA TCT CA AAACT CA TCT CC TCCAA CA | 26 | 3 | -30.669 |
| CA CCACC CA TCT CA AAACT CA TCT CA GCCAA CA | 27 | 6 | -32.377 |
| CA CCACC CA TCT CA AAACT CA TCT CA CCCAA CA | 27 | 5 | -30.706 |
| CA CCACC CA TCT CA AAACT CA TCT CA ACCAA CA | 27 | 4 | -31.367 |
| CA CCACC CA TCT CA AAACT CA TCT CA TCTAA CA | 29 | 8 | -29.403 |
| CA CCACC CA TCT CA AAACT CA TCT CA TCGAA CA | 29 | 9 | -30.705 |
| CA CCACC CA TCT CA AAACT CA TCT CA TCAAA CA | 29 | 7 | -30.378 |

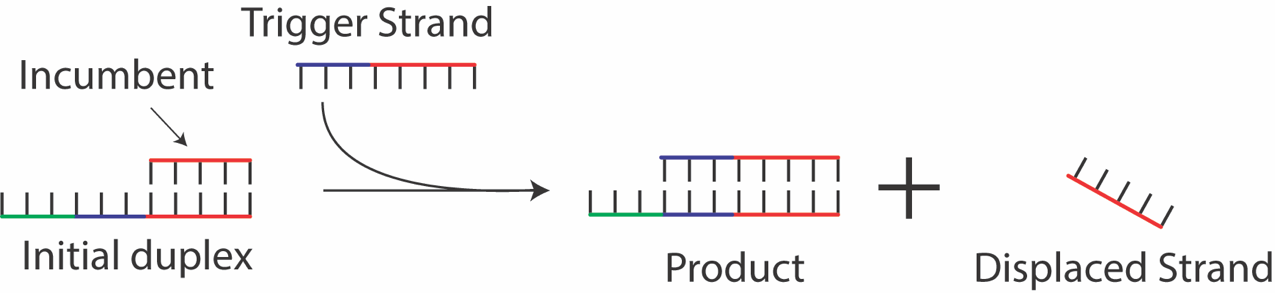

Figure S1: Toehold-mediated strand displacement reaction schematic.

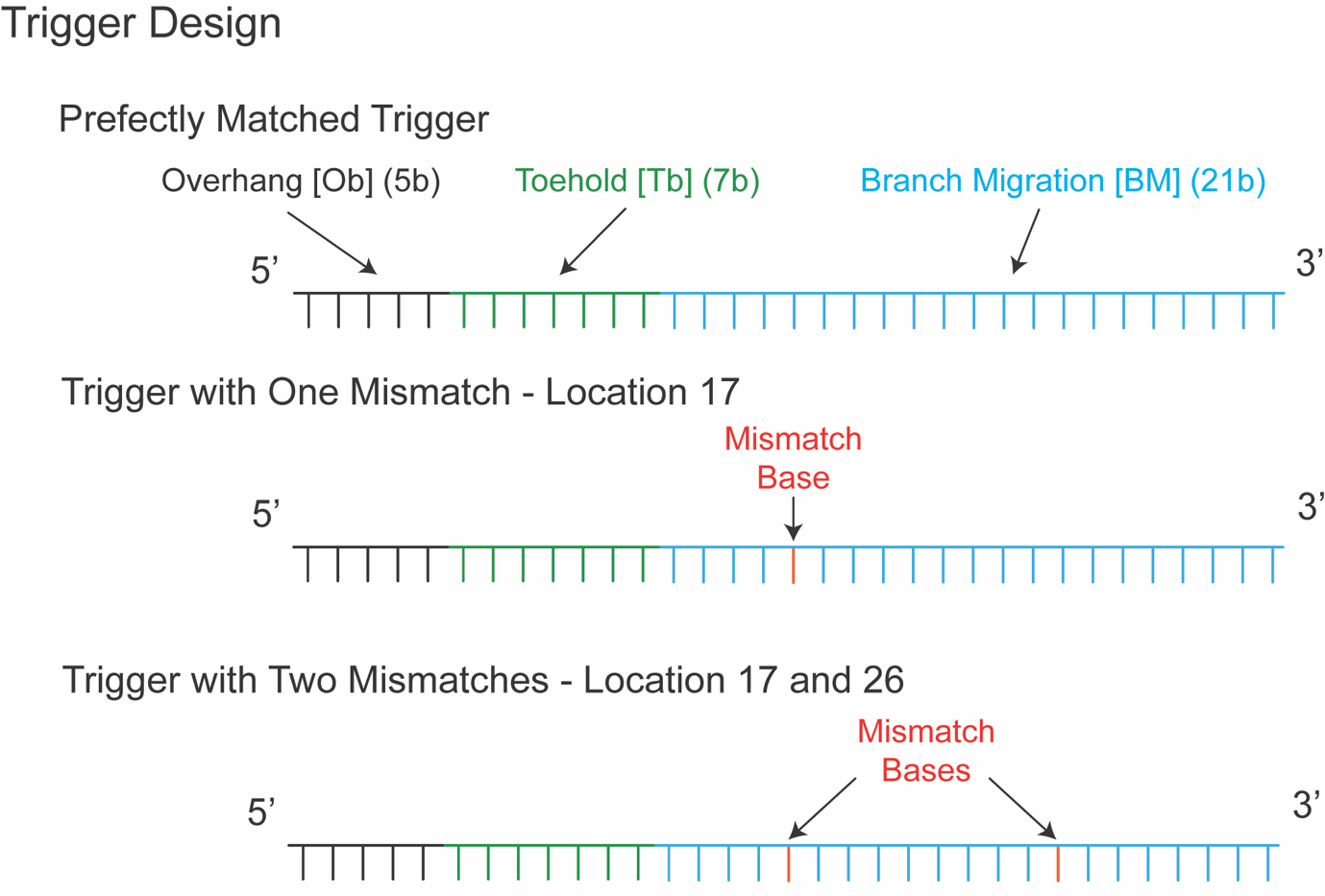

Figure S2: Trigger design schematic showing the three main component of the trigger – overhang, toehold, and branch migration region. The mismatched triggers are shown as the red base(s).

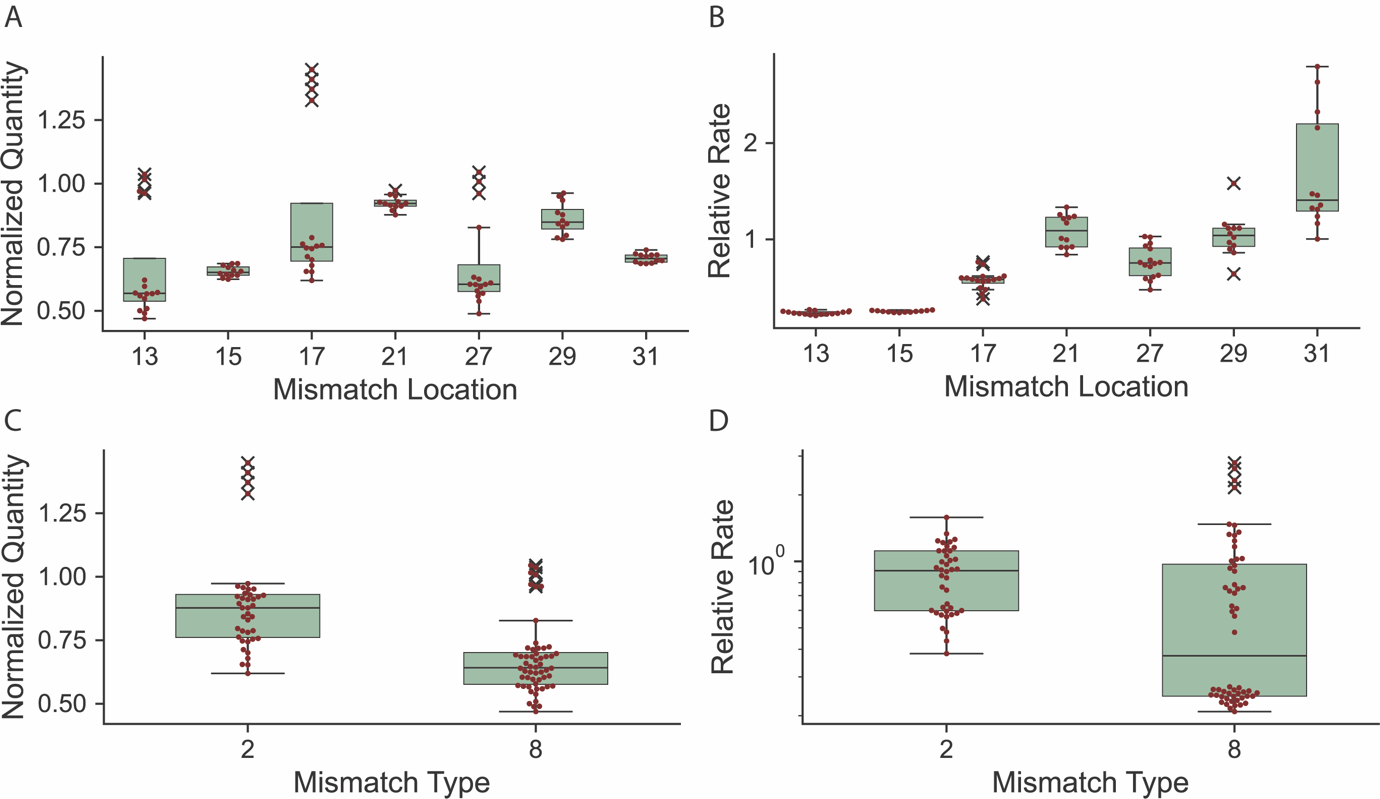

Figure S3: Boxplots of the fitted rates and quantity of triggers of type 2 and type 8 mismatch triggers. (A, B) The fitted final hybridization quantity and rate values based on the mismatch location assuming the two types of mismatches are the same. (C, D) The fitted final hybridization quantity and rate values based on mismatch type.

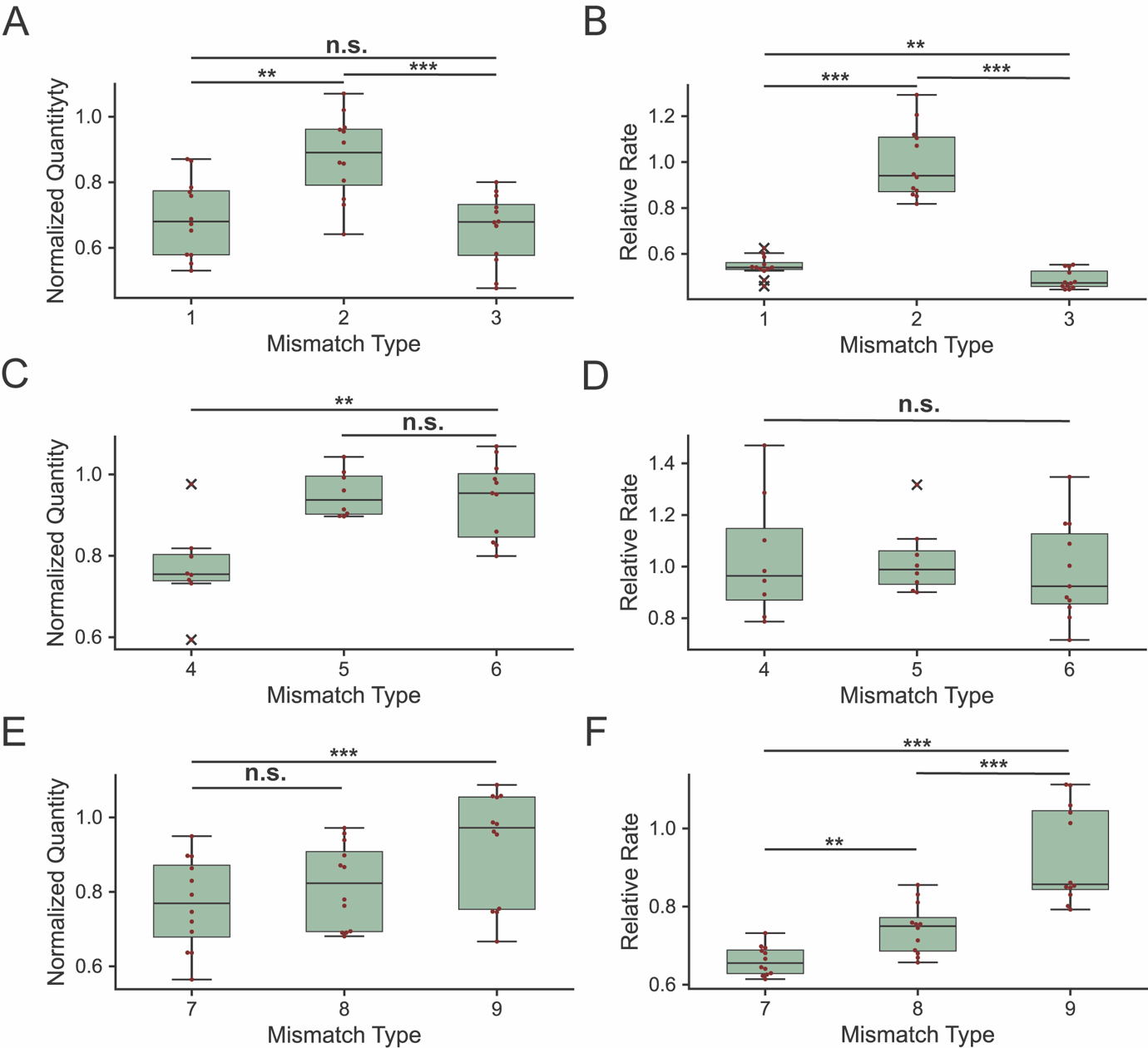

Figure S4: Boxplots of the fitted rates and quantity of triggers of the three mismatch groups using one-phase association model. The trigger was based of T2 shown in Table S1. The mismatch type number corresponds to the number shown in **Table 1** of the main text. (A, B) The fitted final hybridization quantity and rate values for mismatch group 1 at location 17. (C, D) The fitted final hybridization quantity and rate values for mismatch group 2 at location 19. (E, F) The fitted final hybridization quantity and rate values for mismatch group 3 at location 18. Hybridization quantity was normalized against the hybridization quantity of perfectly matched trigger. The relative hybridization was relative to the hybridization rate of perfectly matched trigger. Each dot in the boxplot represents individual measurement. The whisker was set to be the 25^th^ and 75^th^ percentile of the median. The cross represents the potential outlier presented in the dataset (Welch one-way ANOVA with Games-Howell post-hoc analysis, n.s. = no significance, *=p<0.05, **=p<0.01, ***=p<0.001)
